# The ontogeny of multisensory peripersonal space in human infancy: From visual-tactile links to conscious expectations

**DOI:** 10.1101/2020.09.07.279984

**Authors:** Giulia Orioli, Irene Parisi, José L. van Velzen, Andrew J. Bremner

## Abstract

The influence of visual object motion on the processing of bodily events offers a marker for the development of human infants’ perception of themselves in peripersonal space. We presented 4- (n = 20) and 8-month-old (n = 20) infants with an unattended visual object moving towards or away from their body followed by a vibrotactile stimulus on their hands. The 4-month-olds’ somatosensory evoked potentials (SEPs) were modulated by approaching visual motion, demonstrating the early ontogeny of the cortical multisensory foundations of peripersonal space representations. We also observed rapid changes in these markers within the 8-month-old age group: as infants approach 9 months, salient SEP components were increasingly enhanced by (unexpected) tactile stimuli following receding visual motion. These findings provide important clues to the ontogeny of human self-awareness in the first year of life, and suggest important postnatal developments in infants’ expectations about interactions between the body and the external world.

## Introduction

Acting in the environment and comprehending one’s place in it requires an ability to represent the dynamic relationships between sensory events originating in external space (often audiovisual features of objects or people) and those impinging on the body in personal space (often somatosensory inputs)^1–3^. The processing of the spatiotemporal events occurring between external space and the body is referred to as peripersonal spatial representation^4^. The last 20 years have seen significant advances in our understanding of the neural basis of these multisensory representations of peripersonal space^5^ and recent research has also begun to capture the ways in which the human brain makes sensory predictions^6–9^ based on dynamic multisensory interactions at the interface between the body and the world^1,10–12^. However, no research has yet examined the ontogeny of such abilities in early life. Here we trace the early development of dynamic representations of peripersonal space, through an investigation into the influences of visual motion (towards or away from the body) on the processing of subsequently presented somatosensory stimuli, in 4- and 8-month-old human infants. Doing this, we hope to shed light on the emergence in humans of an ability to comprehend the interface between the body and the world.

Over the years there have been a number of attempts to trace the origins of human infants’ ability to perceive what would now be considered as peripersonal spatial events. Since the 1970s, a number of studies investigated young infants’ behavioural responses to visual looming stimuli, focusing on their defensive reactions, especially eye blinks^13–16^. Yonas and colleagues^16^ in particular demonstrated that eye-blinks in response to looming visual events in peripersonal space undergo an extended developmental course, being absent before 2 months of age and reliably present only after 8 months. Schmuckler and colleagues have since demonstrated that the sensitivity of infants’ eye-blinks to looming objects also takes into account the path of approach, and the type of imminent contact^17^. Given the absence of blinking reactions before 4 months of age, Yonas et al.^16^ concluded that young infants are unable to perceive whether an object is approaching their body. More recently however, Orioli et al.^18,19^ demonstrated that when infants’ looking behaviours are measured, even newborns demonstrate an ability to differentiate visual events based on their motion direction relative to the body, showing a visual preference for objects moving towards them vs away from them. As such there may be a developmental offset between perceiving objects moving in peripersonal space and showing defensive reactions to such objects^18,19^. Other studies investigated the perception of multisensory events in peripersonal space, demonstrating that from an early age human infants are sensitive to temporal and spatial multisensory contingencies between visual, auditory and tactile stimuli that are likely to play a fundamental role in peripersonal space representations^20–26^. What remains unclear is the extent to which sensitivity to such multisensory contingencies can support infants’ ability to create spatiotemporally coherent links between visual information specifying motion towards them and subsequent tactile stimulation on the body and, eventually, their ability to predict tactile bodily events.

Research into predictive processing mechanisms in infancy has gained traction in recent years^27–32^, with the broad aim of determining if predictive processing is a continuous ability across the lifespan^32^, as an intrinsic property of the cerebral cortices, or rather if it develops with experience and brain maturation. It appears that 6-month-old infants already show some neural signatures of crossmodal expectation-based feedback across cortical regions^27^ and that 1-year-olds can learn associations between cross-modal events and form predictions and expectations that can influence their neural responses to unexpected or contradictory events^31^. These findings suggest that infants’ early sensory processing could be modulated by top-down influences, supporting the hypothesis that infants can generate an internal model of the environment and form predictions about it^30,33^. According to this framework, the development of infants’ understanding of the physical world could be conceived as the formation of predictive models about the relation between entities in the environment and the infants’ own body and its actions^33^. This may include also infants’ developing ability to perceive, understand and eventually predict the continuity between visual stimuli moving in peripersonal space and tactile stimuli on the body.

Recent accounts of mature representations of peripersonal space in adults have emphasized the fundamental role of inferential and predictive mechanisms in these representational abilities. More specifically, it has been suggested that the special status of the representations of events in peripersonal space (e.g., as observed through speeded responses to objects close to the body) may be explained by the predictive mechanisms at play when somatosensory processing is modulated by prior visual, auditory or audiovisual stimulus perceived near the body and moving towards it^1,12,34^. Correspondingly, a number of studies recently investigated whether responses to tactile stimuli can be modulated by predictive but not spatially or temporally proximal stimuli presented in a different modality^10,11^, and determined that they can. The key novelty of these findings compared with previous research is their focus on crossmodal interactions via predictive relations between visual and somatosensory events, which cannot be mediated via exogenous crossmodal effects due to colocation or synchrony between the visual and tactile stimuli^35,36^. Results showed that the participants detected more easily and responded significantly faster to a tactile stimulus that was presented at the expected time to contact (vs earlier or later) and at the congruent (vs incongruent) location of contact suggested by a visual approaching (but not receding) stimulus^10,11^. These findings support the existence of a predictive mechanism that uses visual motion cues to make judgments about the time and space of an impending tactile stimulus and enhances tactile processing at the time and location of impending contact^11^. However, to our knowledge, no study has, so far, investigated the development of such a mechanism in human infants.

Here, we aimed to address this important gap in our understanding of human development, investigating infants’ ability to perceive multisensory connections between events taking place in peripersonal space and their temporally and spatially separated tactile consequences on the body. We also wanted to explore how, through such multisensory processes, developing humans can build the predictive links allowing them to predict tactile events on the body based on prior sensory events in the environment. To this end, we investigated infants’ perception of continuity between approaching visual stimuli in peripersonal space and tactile stimuli on the body, developing a paradigm similar to that implemented with adults by Kandula et al.^11^.

We recorded the electrical brain activity of a group of 4- and 8-month-old infants presented with tactile stimuli on their hands that were preceded by the visual presentation of an unattended moving object. The two age groups were chosen in light of the several developmental changes taking place between 4 and 8 months of life. For example, infants’ ability to localise touch in relation to external spatial coordinates has been shown to develop between 4 and 6 months^37^ and, relatedly, studies showed that postural information begins to influence the neural correlates of infants’ tactile perception after 6.5 months of life^38^. Additionally, infants’ ability to reach for and handle objects begins to be reliably present around 5 months of life^39–42^. Given the intrinsic link between the ability to act on the environment and to perceive body-related motion in the environment, mastering reaching could have an impact on infants’ tactile prediction. Choosing to include in the present studies infants aged 4 and 8 months allowed us to investigate infants’ prediction of tactile stimuli in the context of these ongoing developmental changes in the first year of life.

To investigate the influence of visual object motion on the processing of bodily events, we presented infant participants with tactile stimuli on their hands preceded by dynamic visual objects on a screen, rendered to specify 3D trajectories either approaching their hands or moving away from them. Our aim was to investigate the predictive influence of the direction of visual motion stimuli that the infants were not visually tracking. To achieve this, an attractive “attention-getter” was presented on the top of the screen throughout the study, and we ensured that the infants focused their gaze on the attention-getter, and not the moving visual stimulus. The approaching display showed a small ball approaching the participants’ hands and disappearing halfway through its trajectory from its starting point to the infant’s hands; the receding display was the approaching sequence of events played backwards. After an interval, which ensured that there was no spatial nor temporal proximity between the visual and the tactile stimulus, the infants felt a tactile stimulus on both hands, which were kept close to each other and held at the expected location of contact as signalled by the approaching visual motion trajectory. The tactile stimulus was presented on 50% of the trials only: in order to measure and compare purely the somatosensory responses, we calculated a difference waveform between the trials where the infants did and did not receive a tactile stimulation. This ensured that the visual components, common across the trials in which the touch was and was not presented, were removed by the subtraction, leaving only the somatosensory evoked potentials (SEPs). We then compared these between the approaching vs receding visual motion conditions.

Given the high adaptive value of perceiving and predicting the contact of visual objects with the body, it would be reasonable to expect that the mechanisms supporting it develop early in life^24^. At the same time, we believe that multisensory postnatal experience would most likely play an important role in infants’ integration of stimuli approaching the body and subsequent tactile stimuli taking place at the expected time and location of contact. This is particularly relevant with regards to visual moving stimuli, which infants would only be able to experience in their postnatal life, contrarily to auditory stimuli. For these reasons, and also in light of the important developmental changes taking place during the first year of life and mentioned earlier^37–42^, we expected to find evidence of developmental changes in the visual modulation of touch in the two age groups who participated in this study. A recent study investigating how infants’ brain responses are influenced by their prior expectations and violations of expectations showed, in 1-year-old infants, an enhancement of early perceptual components in response to predicted vs surprising cross-modal events^31^. In light of this result, we expected that if infants could perceive the continuity between approaching motion and subsequent tactile stimuli on the body, they would show larger SEPs in response to the tactile stimulus following approaching vs receding motion. This hypothesised pattern of results would also be consistent with recent findings in adults where, when participants anticipated a tickling sensation, enhanced activity in contralateral primary somatosensory cortex was observed^43^.

## Results

We recorded infants’ brain responses to vibrotactile stimuli on the palms of the hands preceded by unattended visual stimuli that either loomed towards (Approaching) or moved away (Receding) from the infant’s hands. We aimed to provide a valid comparison of the effects of approaching and receding visual stimuli on somatosensory processing, unpolluted by differences in the response determined by the visual components of the stimulation. In order to do so, we computed the somatosensory evoked potentials for vibrotactile events in the Approaching and Receding conditions, and then subtracted from these the responses recorded in “No-Touch” trials, where only the visual approaching and receding stimuli were presented. This step ensured that any differences in the SEPs measured on the scalp were not determined by the infants’ brain responses to the visual stimuli alone. For both age groups, we analysed the SEP responses originating from sites close to somatosensory areas, in the regions surrounding CP3 and CP4 in the 10-20 system. Because ERP components change significantly in amplitude and latency during the first year of life^44,45^, we treated the responses of the 4- and the 8-month-old infants separately.

### 4-month-olds

First, in order to trace the time course of statistically reliable modulations of the SEPs by Condition, we ran a sample-point by sample-point analysis, using the Monte Carlo simulation method^46^, controlling for autocorrelation between sample points. Supplied with the individual averaged amplitude of the difference waveforms between 100 ms prior to the tactile stimulus onset and 900 ms after it, the simulation identified as reliably significant any sequence of consecutive significant t-tests longer than 220 ms (estimated auto-correlation at lag 5 = 0.986) and highlighted a reliably significant sequence between 202 and 700 ms after the onset of the tactile stimulus (Fig. 1A). Within this time window we identified, by visual inspection, five components on which we focused for further analyses [P286 (202-354 ms); N398 (356-440 ms); P506 (442-548 ms); N560 (550-598 ms); P662 (600-700 ms)]. In order to avoid biased measurements, the exact time range for each component was determined using a collapsed localisers approach^47^: we averaged the data across participants and conditions and then selected the time range showing the largest activity for each component. We then used this time range to measure the mean individual amplitude of the response in each component for the two conditions separately (Fig. 1B). Paired planned comparisons confirmed that, across all components, the amplitude of the response was always significantly larger when the tactile stimulus had been preceded by approaching rather than receding motion (see Table 1).

**Figure 1.**
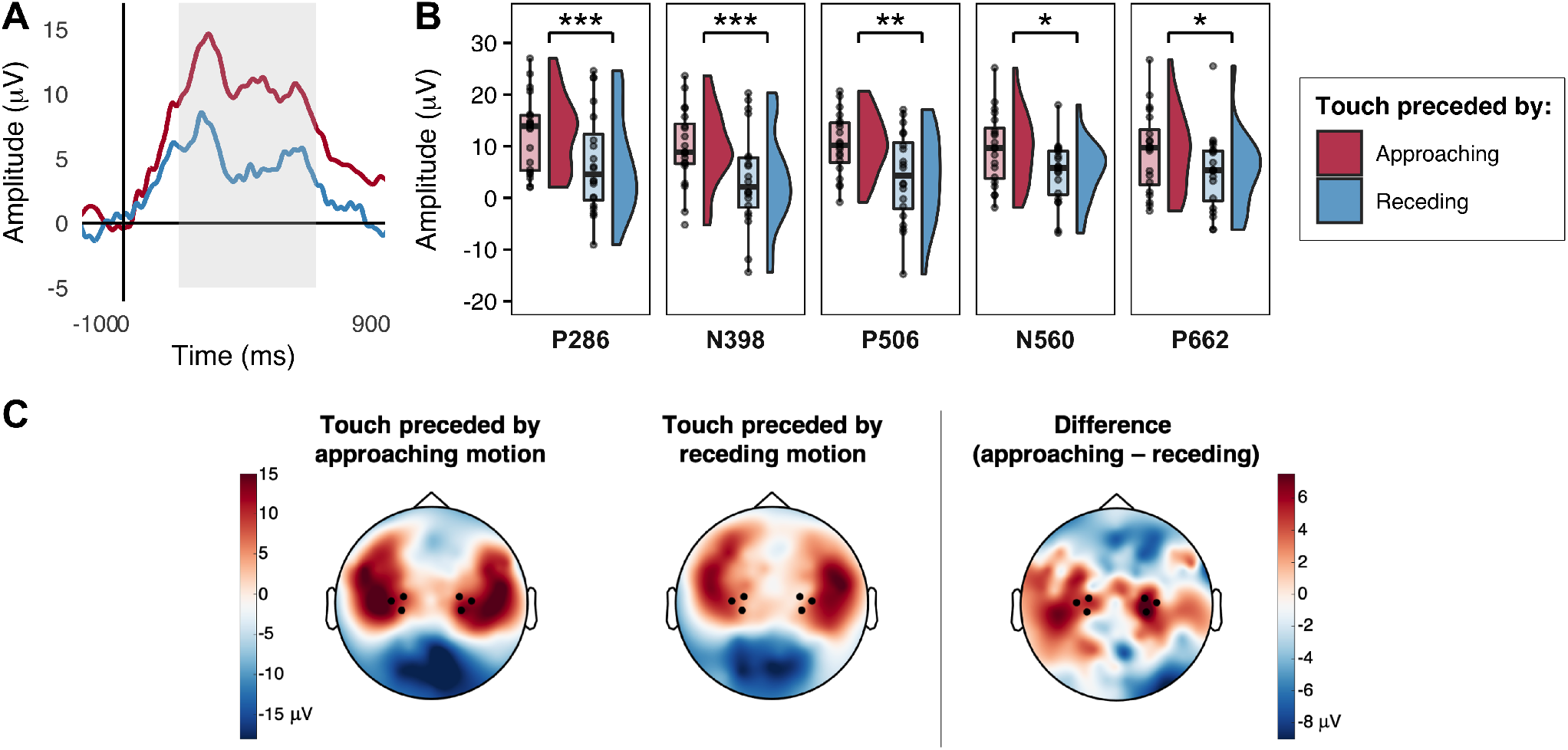
Modulation of 4-month-old infants’ SEPs by the direction of prior unattended visual motion (approaching vs receding). **A**. Grand average SEPs across hemispheres; the reliably significant difference identified by the sample-point by sample-point analysis is indicated by the grey shading. **B**. Voltage differences in the grand averaged mean individual amplitude of the SEPs in the two conditions for the five components of interest; significant comparisons are indicated (* = *p* < 0.05, ** = *p* < 0.01, *** = *p* ≤ 0.001). **C**. Grand average topographical representations of the voltage distribution over the scalp in the two conditions between 202 and 700 ms after the tactile stimulus onset (the period of time during which the sample-point by sample-point analysis revealed a statistically reliable difference), with a Touch following Approaching motion - Touch following Receding motion difference map to the right; channels averaged for the analyses are highlighted.

**Table 1.** Results of the paired planned comparisons on the mean individual average of the amplitude of the SEPs in Approaching vs Receding conditions for each component occurring between 202 and 700 ms post stimulus onset (including Kolmogorov-Smirnov tests, *D,* for testing the normality of the distribution of the differences between conditions).

| **Component** | ***D value*** | ***p value*** | ***dfs*** | ***t value*** | ***p value*** | ***d<sub>z</sub>*** |
| --- | --- | --- | --- | --- | --- | --- |
| P286 | 0.111 | 0.945 | 19 | 3.717 | 0.001 | 0.831 |
| N398 | 0.165 | 0.591 | 19 | 3.803 | 0.001 | 0.850 |
| P446 | 0.119 | 0.909 | 19 | 3.463 | 0.003 | 0.774 |
| N560 | 0.157 | 0.652 | 19 | 2.518 | 0.021 | 0.563 |
| P622 | 0.115 | 0.928 | 19 | 2.490 | 0.022 | 0.557 |

### 8-month-olds

As for the younger age group, we ran a sample-point by sample-point analysis, using the same Monte Carlo simulation method^46^. The simulation identified as reliably significant any sequence of consecutive significant t-tests longer than 106 ms (estimated auto-correlation at lag 5 = 0.867). Based on this criterion, the analysis did not identify any sequences of sample points that were reliably different between conditions (Fig. 2A). Given that this method is insensitive to differences that occur only on brief segments of time^46^, we further probed those components that were identifiable by visual inspection within the time window of differences highlighted by the sample-point by sample-point analysis in the 4-month-olds group (i.e., between 202 and 700 ms after the onset of the tactile stimulus). Four components were identified, using the same method used for the 4-month-olds’ group^47^ [P240 (202-310 ms); N362 (312-418 ms); P470 (420-526 ms); N572 (528-636 ms)]. Planned comparisons on the mean individual amplitude of the response between conditions in each component revealed no significant differences, confirming the results of the sample-point by sample-point analysis (see Fig. 2B and Table 2).

**Figure 2.**
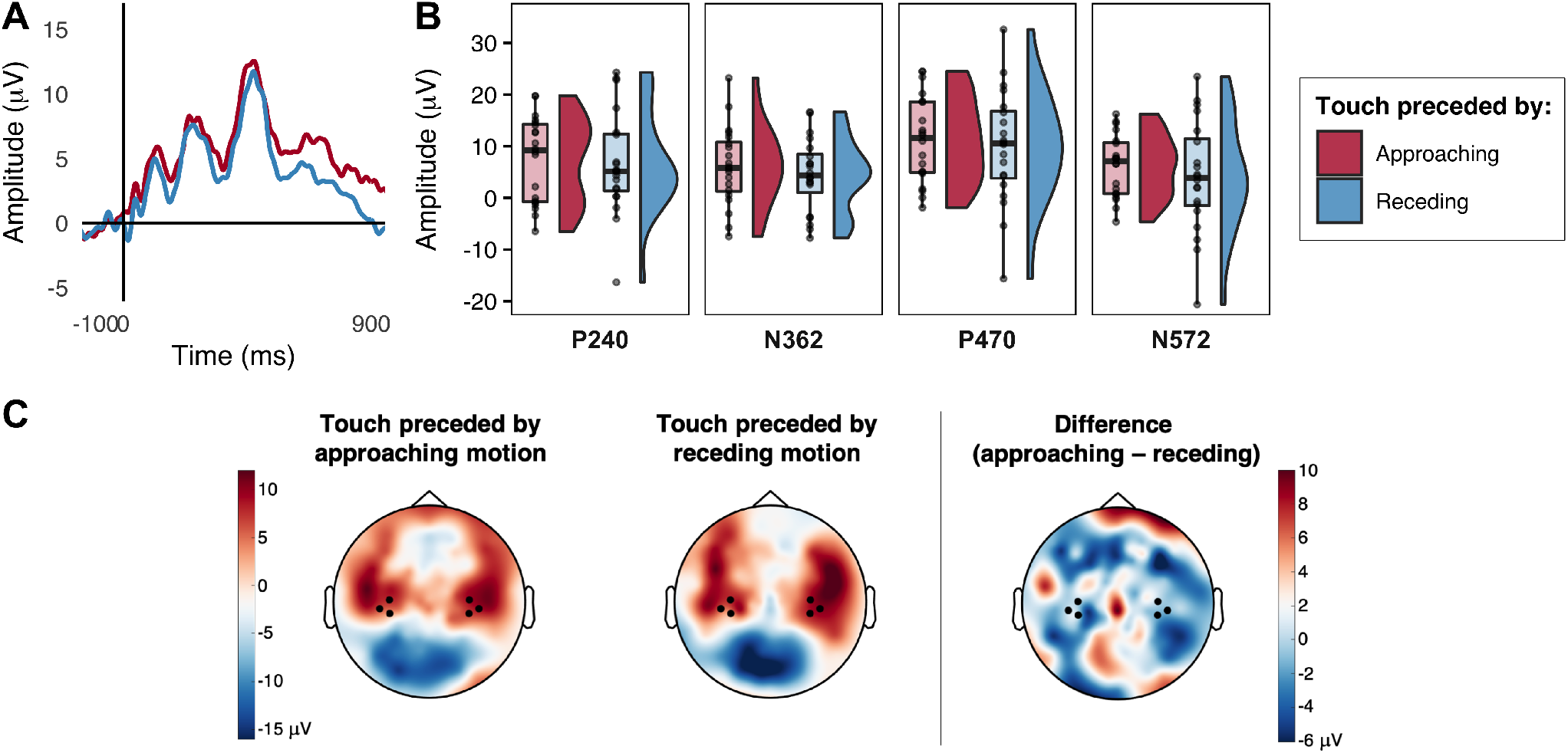
Modulation of 8-month-old infants’ SEPs by the direction of prior unattended visual motion (Approaching vs Receding). **A.** Grand average SEPs across hemispheres. **B.** Voltage differences in the grand averaged mean individual amplitude of the SEPs in the two conditions for the four components of interest. **C.** Grand average topographical representations of the voltage distribution over the scalp in the two conditions between 202 and 700 ms after the tactile stimulus onset (the period of time during which the sample-point by sample-point analysis revealed a statistically reliable difference in the 4-month-olds group), with a Touch following Approaching motion - Touch following Receding motion difference map to the right; channels averaged for the analyses are highlighted.

**Table 2.** Results of the paired planned comparisons on the mean individual average of the amplitude of the SEPs in Approaching vs Receding conditions for each component occurring between 202 and 700 ms post stimulus onset (including Kolmogorov-Smirnov tests, *D*, for testing the normality of the distribution of the differences between conditions).

| Component | *D value* | *p value* | *dfs* | *t value* | *p value* | *d<sub>z</sub>* |
| --- | --- | --- | --- | --- | --- | --- |
| P240 | 0.194 | 0.387 | 19 | 0.490 | 0.630 | 0.109 |
| N362 | 0.133 | 0.828 | 19 | 0.867 | 0.397 | 0.194 |
| P470 | 0.118 | 0.913 | 19 | 0.599 | 0.556 | 0.134 |
| N572 | 0.122 | 0.894 | 19 | 0.937 | 0.361 | 0.210 |

The absence of any effect of the direction of prior unattended visual motion on somatosensory processing in 8-month-old infants is surprising in the context of the robust effect observed in 4-month-olds. Given the need to better understand this surprising developmental change, we decided to explore the 8-month-olds data in more detail. Specifically, we investigated whether any differences between conditions could have been masked by individual variations between participants, such as their developmental status. Using age in days as a proxy for precise developmental status within the 8-month-old age group, we fitted 3 linear mixed-effect models^48^ including a categorical fixed effect (Condition), a continuous fixed effect (Age in days) and a random effect (the individual Participant), to explain variation in each of the four main components identified above (P240, N362, P470, N572). The first model (*m1*) included only Condition as a fixed effect and Participant as a random effect (replicating the factors included in the previous analyses). The second model (*m2*) added Age as a second fixed effect. Finally, the third model (*m3*) added the Interaction between Condition and Age. The assumptions of linearity, homoskedasticity and normal distribution of the residuals and the random effects were met by all models for each component.

Likelihood Ratio Tests (LRTs) were conducted to compare how well the three models explained the data. In the first 3 components (P240, N362 and P470), *m3* explained the collected data better than any of the other models. In the N362 component, also *m2* explained the data better than *m1*, while this was not the case for the two positive components. In the fourth component (N572), *m2* was the best fit model (the results of the LRTs between the models are summarised in Table 3).

**Table 3.** Results of the LRTs comparing the 3 models used to analyse the effects of Condition, Age (in days) and their Interaction on the SEPs of 8-month-old infants; significant comparisons are indicated (* = *p* < .05, ** = *p* < .01).

|  | ***m2 vs m1*** |  |  |  | ***m3 vs m1*** |  |  |  | ***m3 vs m2*** |  |  |  |
| --- | --- | --- | --- | --- | --- | --- | --- | --- | --- | --- | --- | --- |
| | *df* | $\chi^2$ | *p-value* | | *df* | $\chi^2$ | *p-value* | | *df* | $\chi^2$ | *p-value* | |
| **P240** | 1 | 0.677 | 0.411 |  | 2 | 7.317 | 0.026 | * |  |  |  |  |
| **N362** | 1 | 7.598 | 0.006 | ** |  |  |  |  | 1 | 7.739 | 0.005 | ** |
| **P470** | 1 | 2.876 | 0.090 |  | 2 | 7.390 | 0.025 | * |  |  |  |  |
| **N572** | 1 | 4.822 | 0.028 | * |  |  |  |  | 1 | 0.246 | 0.620 |  |

We deal first with the findings regarding the first 3 components (P240, N362 and P470) where *m3* was the best fit. The *m3* model is the only one that included the interaction between Condition and Age. This term was significant across all three components [P240: *t*(18) = 2.662, *p* = 0.016; N362: *t*(18) = 2.916, *p* = 0.009; P470: *t*(18) = 2.135, *p* = 0.047]. This showed how, with increasing age in days, infants SEPs changed from demonstrating an enhanced response to tactile stimuli preceded by approaching visual motion to demonstrating an enhanced response to tactile stimuli preceded by receding visual motion (see Fig. 3A). The results also highlighted a significant main effect of Condition on these components once Age and the random effect of Participants were taken into account [P240: *t*(18) = −2.679, *p* = 0.015; N362: *t*(18) = −2.947, *p* = 0.009; P470: *t*(18) = −2.155, *p* = 0.045]. In the fourth component (N572), *m2*, including the fixed effects of Condition and Age, and the random effect of participant, was the best fit. Reflecting the better fit compared to *m1* there was a main effect of Age on the amplitude of the N572 [*t*(18) = −2.215, *p* = 0.04]: this describes a decline in the amplitude of the SEPs with age across both conditions (Fig. 3A).

**Figure 3.**
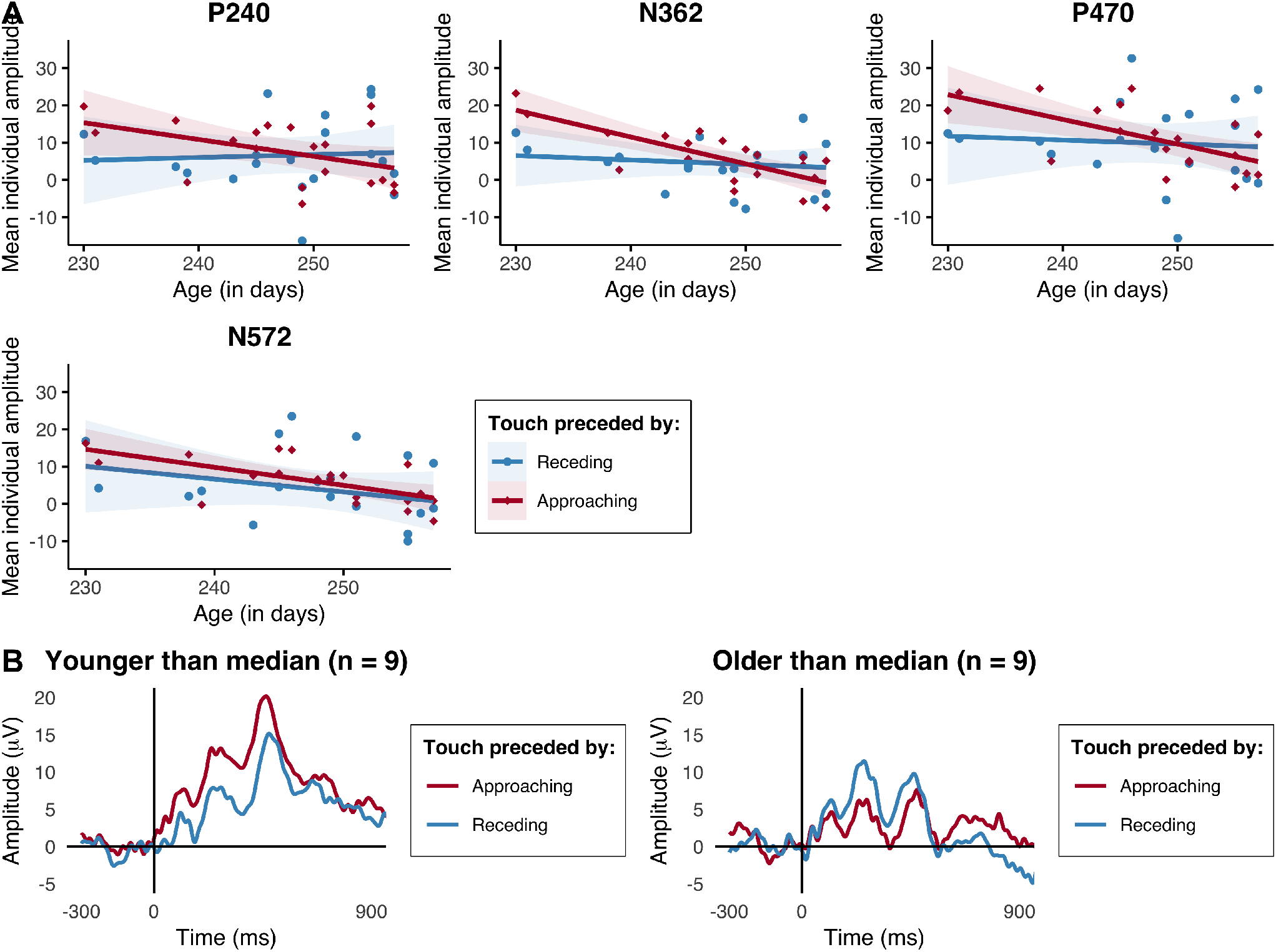
Effect of Condition and Age (in days) on the SEPs of 8-month-old infants. **A.** Scatter plots illustrating, for each component of interest, the relationship between the mean individual amplitude of the SEPs in each condition and the infants’ age in days, with regression lines (and S.E., shaded) for each condition. **B.** For illustrative purposes, in light of the results of the LMMs, we plotted the grand averaged SEPs for the younger and the older 8-month-olds (2 groups of 9 infants each, created based on the median age value, 249 days); the plots suggest that the direction of the difference between the SEPs in response to a tactile stimulus following approaching vs receding motion might reverse between the younger and the older 8-month-old infants.

Altogether these results indicate that the amplitude of the salient SEP components in 8-month-olds can be modulated by whether prior to the tactile stimulus they perceive a visual object approaching them or receding away from them. This is observed via an interaction between the effect of the visual condition they were presented with (approaching vs receding motion) and their age in days. More specifically, younger 8-month-old participants show, as 4-month-olds, a larger response to tactile stimuli preceded by approaching motion, while older 8-month-old participants show the opposite pattern (Fig. 3B).

For completeness, we fitted the same models for the 4-month-old infants. In this group, neither *m2* nor *m3* significantly improved on the fit of *m1*, which included only Condition as a fixed effect and Participant as a random effect. This was the case for all of the five components tested (the results of the LRTs between the models are summarised in Table 4). This result confirms our previous findings that 4-month-old infants demonstrate an enhancement of the response to the tactile stimulus when it was preceded by unattended approaching vs receding visual motion, throughout all five components analysed and irrespective of the participants’ precise ages in days (Fig. 4).

**Figure 4.**
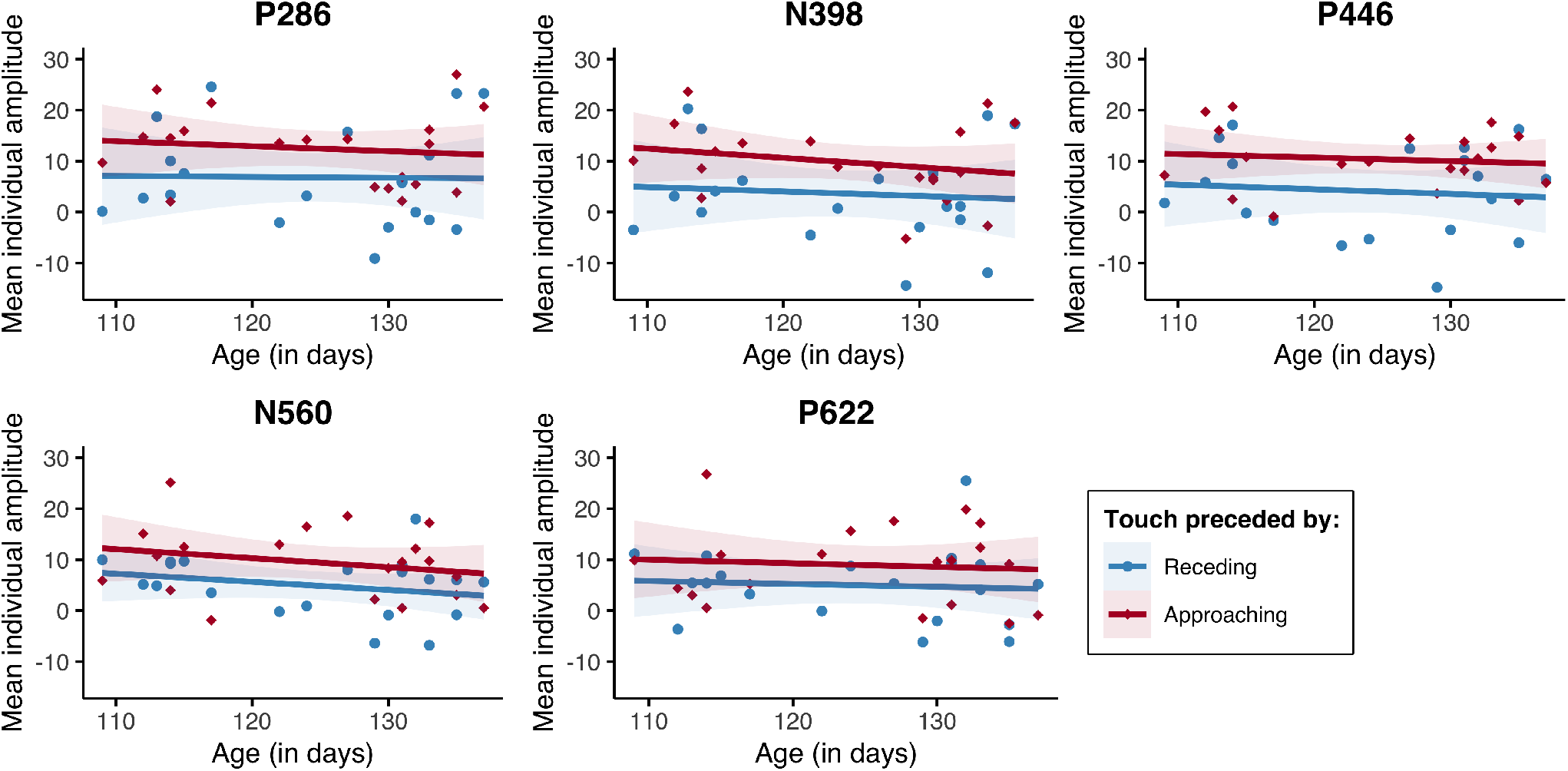
Effect of Condition and Age (in days) on the SEPs of 4-month-old infants. Scatter plots illustrating, for each component of interest, the relationship between the mean individual amplitude of the SEPs in each condition and the infants’ age in days, with regression lines (and S.E., shaded) for each condition.

**Table 4.** Results of the LRTs comparing the 3 models used to analyse the effects of Condition, Age (in days) and their Interaction on the SEPs of 4-month-old infants.

|  | *m2 vs m1* |  |  | *m3 vs m1* |  |  |
| --- | --- | --- | --- | --- | --- | --- |
| | *df* | $\chi^2$ | *p-value* | *df* | $\chi^2$ | *p-value* |
| **P286** | 1 | 0.085 | 0.770 | 2 | 0.338 | 0.844 |
| **N398** | 1 | 0.534 | 0.465 | 2 | 0.825 | 0.662 |
| **P506** | 1 | 0.285 | 0.594 | 2 | 0.296 | 0.862 |
| **N560** | 1 | 1.949 | 0.163 | 2 | 1.957 | 0.376 |
| **P662** | 1 | 0.153 | 0.696 | 2 | 0.160 | 0.923 |

## Discussion

Here we show, for the first time, that human infants from at least 4 months of age represent multisensory predictive interactions between visual and tactile events across peripersonal space. In the study reported here we measured 4- and 8-month-old infants’ SEPs following the presentation of unattended visual objects that either approached or receded from the infants’ own bodies (their hands). We found that 4-month-olds’ (and younger 8-month-olds’) SEPs are significantly enhanced following approaching visual motion, while older 8-month-olds seem to show the reverse pattern, with larger SEPs in response to tactile stimuli following receding motion. This indicates that unattended visual information specifying the motion of an object in relation to an infant observer in peripersonal space (towards or away from their bodies) affects subsequent somatosensory processing in both 4- and 8-month-old infants.

A number of previous findings showed that infants, from as early as birth, are able to perceive the direction of visual and audiovisual motion towards or away from themselves^18,19,23^. However, here we establish for the first time that sensitivity to cues concerning the movements of external objects in peripersonal space also impacts on infants’ representations of tactile stimuli on their own bodies^49^. The ability to perceive oneself as situated in the external environment is underpinned by just such multisensory interactions between the senses specifying external objects and events (e.g., vision and hearing) and those more directly specifying bodily perception (touch and vestibular balance). The present findings therefore prompt the striking conclusion that human infants, from as young as 4 months of age, are endowed with some of the key multisensory abilities that enable mature humans and animals to sense their bodily selves in relation to the visual external world that they inhabit.

The fact that we have established these abilities in 4-month-old infants is significant. Infants do not typically make successful reaches to objects that they have targeted in vision before 5 months of age^39–42^, and so our results suggest that we should entertain the possibility that infants can learn about these visual-tactile links irrespective of an ability to undertake skilled action in the external environment. Indeed, it seems likely that infants will be exposed to rich visual-tactile multisensory experiences in the first postnatal months (e.g., in the context of infant-parent tactile interactions such as breastfeeding and tickling), which could provide the basis for learning about the associations between visual motion and tactile contact that are necessary to explain our findings. Further research investigating the nature of early multisensory experiences in the first months of life^50^ could shed valuable light on the early origins of infants’ perceptions of the links between their bodies and the visual environment.

Importantly, the crossmodal links between visual and tactile cues that we showed here are not explicable through straightforward temporal and/or spatial coherence^22,51^: in the present study, visual cues specifying approaching or receding motion affected subsequent processing of a tactile stimulus presented in a different place in external space and at a different point in time, after the visual cue had disappeared. However, whilst we have demonstrated that infants’ somatosensory systems can be modulated by visual information presented at a prior moment in time and at a distant location in space, there are a number of means by which this might be achieved. For instance, it may be possible to explain such abilities via infants’ extended crossmodal temporal binding windows^52^. Previous research demonstrated developmental narrowing of the visual-tactile temporal binding window in childhood, showing that 7-year-old children are more likely to consider simultaneous a visual and a tactile event separated by more than 200 ms^52^. We might extrapolate that there is narrowing of such binding windows also between early infancy and later development, and therefore estimate infants’ visual-tactile window to be even longer, while the temporal gap between the end of the visual stimulation and the beginning of the tactile stimulation was only 333.3 ms long. By extension, it may be that the infants in our study showed visual modulations of somatosensory processing by virtue of a simultaneous perception of the visual and tactile events, rather than via a somatosensory expectation based on the direction of the visual motion. If this were the case, 4-month-old infants’ larger responses to the tactile stimuli following approaching motion could be related to younger infant’s behavioural preferences for congruent crossmodal stimuli^25^, wherein congruency may be defined as something that frequently happens together in their everyday life experience. Older 8-month-old infants’ larger responses to tactile stimuli following receding motion, instead, could be linked to older infants’ behavioural preferences for incongruent crossmodal stimuli, which contradict their experience^22,25^.

In line with a number of other studies concerning the development of body representations and somatosensory processing in early life^37,38,53^, we have uncovered evidence of considerable developmental change in the visual modulation of somatosensory processing between 4 and 8 months of age. The enhancement of somatosensory responses seen in 4-month-olds when tactile stimuli are preceded by unattended visual cues specifying approaching object motion is no longer apparent in 8-month-olds. We might attribute this to a better attentional control in the older 8-month-old infants, leading them to focus more on the attention getter and to completely ignore the peripheral visual motion. However, we believe this explanation is unlikely based on previous findings demonstrating behavioural facilitation effects of unattended approaching visual motion on tactile detection in adults, whose attentional control is more efficient than infants’^10,11^. Further analyses using 8-month-old infants’ age in days to predict differences in SEPs across condition showed that the enhancement of somatosensory processing by prior unattended visual approaching motion is present in the younger 8-month-olds, but gradually reverses such that, by 260 days of age, infants are showing enhanced SEPs on prominent components (P240, N362 and P470) following prior visual cues specifying receding motion. This reversal across a short developmental time span is intriguing. One possible explanation for the developmental emergence of enhanced responses to tactile stimuli preceded by receding motion is that this represents the development of a neural process involved in signalling prediction error (i.e., signalling that the tactile contact experienced following receding motion is unexpected). As such, the gradual emergence across the 8-month-old group of greater responses to tactile contact that is not anticipated by prior unattended visual motion information might represent the emergence at this age of top down influences of prediction on the perceptual processing of somatosensory information. This explanation in terms of a developmental emergence of greater responses to unexpected outcomes gains support from a similar developmental change in behavioural responses: younger infants tend to show behavioural (visual looking) preferences for colocated, and synchronous crossmodal stimuli^25^, with older infants tending to look longer at crossmodal relations that contradict typical experience.

Newborn infants are able to distinguish aspects of the visual environment that specify the motion of objects with respect to their field of vision. However, in order to determine whether they perceive the motion of objects with respect to their own selves (their own bodies), a crucial test is whether an unattended visual approaching object might influence how they process bodily events. Here we showed that, from as early as 4 months of age, human infants process somatosensory information differently when it has been preceded by a temporally and spatially distant visual object that approaches the body. This finding indicates that fundamental aspects of the multisensory processes underpinning peripersonal space representations, and self-awareness more generally, are in place prior to the onset of skilled action. Nevertheless, there are striking developmental changes in how infants’ brains process visual-tactile events occurring across peripersonal space between 4 and 8 months of age. As infants approach 9 months we increasingly see, in later somatosensory components, a greater processing of those tactile stimuli that were not predicted by preceding unattended visual motion. These findings yield exciting new clues to the ontogeny of human self-awareness in the first year of life, suggesting important postnatal developments in the ability to form expectations about the interactions between the body and the external environment.

## Method

### Participants

The 4-month-old age-group (n = 20) included 9 female and 11 male infants, with an average age of 124.65 days (*SD* = 9.24 days). A further 31 4-month-olds participated in the study, but were excluded due to fussiness or excessive movement (9), experimental error (3), high impedance around the vertex, which served as reference (1), or due to their having contributed an insufficient number of artefact-free trials in each condition (18). The 8-month-old age-group (n = 20) included 10 female and 10 male infants, with an average age of 247.5 days (*SD* = 8.06 days). A further 34 8-month-olds participated in the study, but were later excluded due to fussiness or excessive movement (14), experimental error (1), high impedance around the vertex (3), or due to their having contributed an insufficient number of artefact-free trials in each condition (16). All of the participants were recruited through the Goldsmiths InfantLab database and received a small gift as a compensation for participating in the research. The testing sessions took place when the infant was awake and alert, ideally at a time that suited their daily routine as advised by the parents. The parents were informed about the procedure and provided Informed Consent to their child’s participation. Ethical approval was gained from the Ethics Committee of the Department of Psychology at Goldsmiths, University of London.

### Design

The study included two conditions, based on the direction of the visual motion events presented to the infants: Approaching and Receding. Each condition comprised two types of trials: Touch and No-Touch. In both types of trials of each conditions, the infants were presented with a set of visual events on a screen (see “Procedure, stimuli and apparatus” section below), comprising an attention-getting stimulus in the top half of the screen, and visual motion events in the bottom half of the screen (approaching or receding depending on the condition). In the Touch trials, but not in the No-Touch trials, these visual events were followed by vibrotactile events on both hands. The No-Touch trials were included in order to make sure that the analysed waveforms were not affected by the visual components of the stimulation. To achieve this, we computed the difference waveforms obtained subtracting, for each visual motion condition (Approaching vs Receding), the response recorded in the No-Touch condition from the response recorded in the Touch condition. The difference waves obtained represented the response to the tactile stimulus itself, given that visual components, common across the two types of trials, were removed by the subtraction. The trials were presented in groups of 4, within which each condition and trial type was presented in a random order, and grouped in blocks of 8 trials each. After each block, the experiment was paused and the infant was presented with short videos (12 s long), intended to break up the repetitiveness of the experimental stimuli and recover their interest and attention. Each infant was presented with a maximum of 10 blocks (20 trials per trial type), until their attention lasted.

### Procedure, stimuli and apparatus

Each infant sat on their parent’s lap on a chair positioned in front of a 24” screen in a dimly lit room. The parent was instructed to keep their child’s hands close to each other about 30 cm away from the screen, and to hold them as still as possible for the duration of the study. At the beginning of the study, an infant-friendly music video was played to attract the infant’s attention to the screen and to help them settle in the experimental room. As the infant was attending to the screen, the experiment began.

Throughout each trial, the infant was presented with an “attention-getting” animal character face located in the top half of the screen. This was constantly rotating in the picture plane, alternately clockwise and anticlockwise (the first direction of motion was counterbalanced across participants), between an orientation where the upper portion of the vertical axis of the animal face was 45° anticlockwise from the environmental vertical, and one in which the upper portion of the vertical axis of the animal faces was 45° clockwise from the environmental vertical. The cartoon animal face subtended a visual angle of 22.17° x 20.43° and was selected randomly on a trial-by-trial basis from a set of 10 possible faces. These attention-getters were intended to attract and hold the participant’s attention for the whole duration of the trial. Any trials in which the infant shifted their gaze away from the attention getter during the trial were identified by offline coding of the infant’s looking behaviour and were excluded from the analyses.

Once the infant was fixating on the attention-getting animal face, the experimenter triggered the presentation of the experimental stimuli. A 3D rendered red ball appeared in the lower half of the screen and either approached the infant’s hands or receded towards the background (approaching and receding events lasted 333.3 ms and comprised 10 images and 20 frames). At its smallest size, the ball subtended a visual angle of 5.90° x 5.57°. The ball moved within a 3D rendered room, whose width, at the bottom of the screen, measured 40 cm. This width was used as a common reference between the real and the simulated worlds in order to calculate the distance of the simulated background wall from the screen surface. We wanted the screen surface to be perceived halfway between the background wall and the infant’s hands, which were located 30 cm away from the screen. Therefore, considering that the 40 cm width of the simulated room corresponded to 28 measurement units in the rendering software, we located the background wall 21 measurement units away from the front of the screen. In this way we obtained a simulated back wall located 30 cm away from the screen surface and, in turn, 60 cm away from the participants hands. Under this rendering, then, the screen surface was specified halfway between the simulated background wall and the participants’ hands. In the Approaching trials, the ball moved from the background wall towards the infant’s hands but disappeared when the rendering specified that it had reached the screen (at bottom of the display). Because the infant’s hands were placed 30 cm away from the screen, the rendering presented the impression the ball disappeared halfway through its trajectory from the background wall to the infant’s hands. In the Receding trials, the simulated motion of the ball specified a trajectory from the halfway point towards the simulated background wall. In both Approaching and Receding trials, after the ball disappeared there was an interval (“gap”) lasting as long as the motion (333.3 ms), following which the infant received, on 50% of the trials, a vibrotactile stimulus on both hands, lasting 200 ms (Fig. 5 and Supplementary Movie 1). Given that the ball disappeared when it reached the simulated halfway point between the background and the infant’s hands, presenting the tactile stimulus after an interval lasting as long as the motion ensured that this was presented at the expected time to contact of the simulated moving ball with the hands in the Approaching condition. Each trial lasted a minimum of 4 s, including minimum 2 s (or as long as the infant needed) when only the attention-getter animal face was presented, followed by 333.3 ms of visual motion and 333.3 ms of gap, and finally by 1.33 s of response collection time, the first 200 ms of which corresponded to the tactile stimulation in the Touch trials.

**Figure 5.**
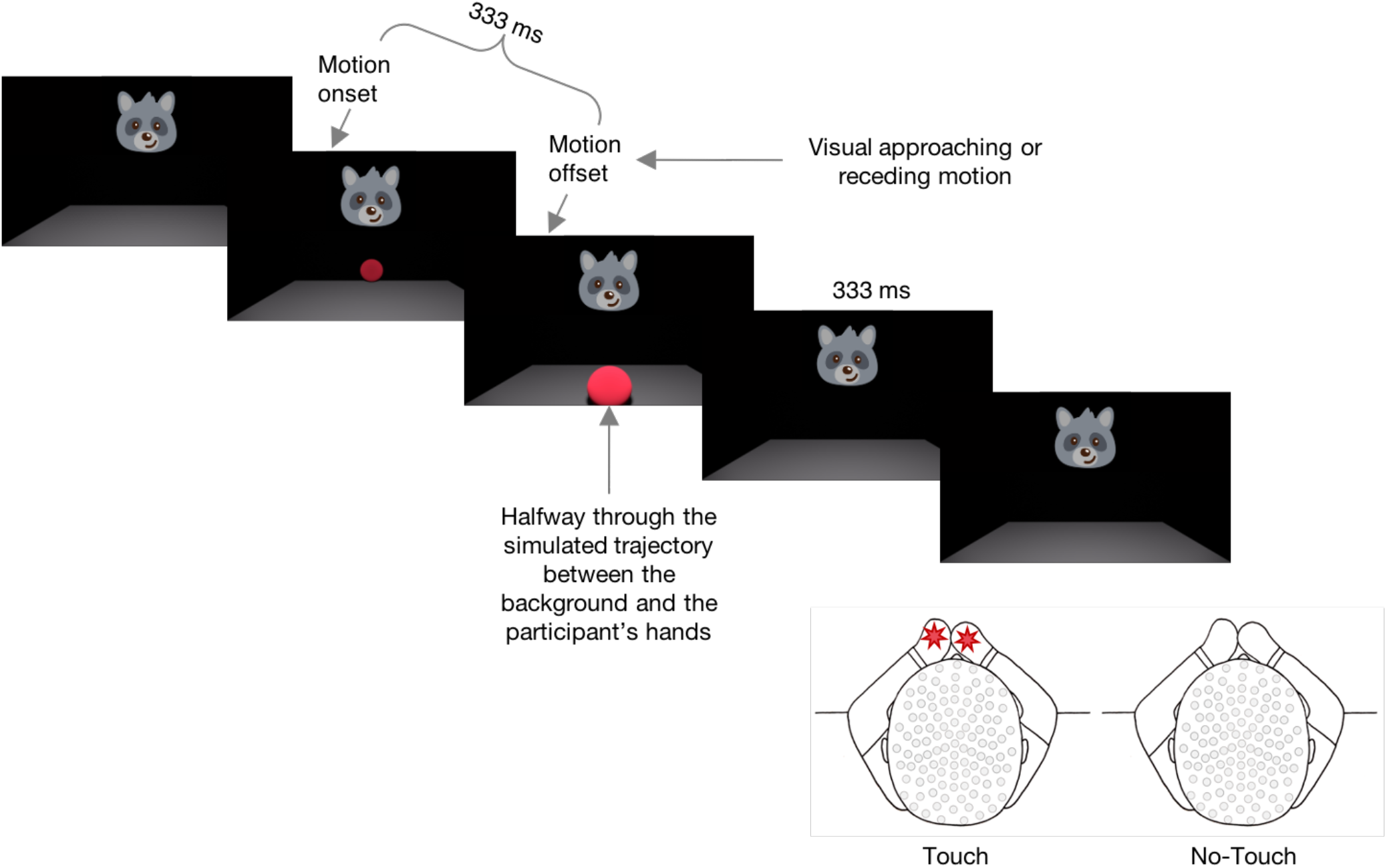
Schematic representation of the Experimental Procedure. When the infant fixated on the attentiongetting animal face, the experimenter triggered the presentation of the experimental stimuli. A red ball appeared in the lower half of the screen and either approached the infant’s hands or receded towards the background for 333.3 ms. After a 333.3 ms interval, the infant received, on 50% of the trials, a vibrotactile stimulus on both hands, lasting 200 ms. During the study, the parent was instructed to keep the infant’s hands close to each other and along the midline, i.e. along the simulated trajectory of the moving ball.

The vibrotactile stimuli were delivered via custom-built voice coil tactile stimulators (tactors), driven by a 220 Hz sine wave. One tactor was placed in each of the infant’s hands and fixed to the palms with self-adherent bandage; the infant’s hands and the tactors were then covered with small cotton mittens. In order to mask the noise of the tactors, an audio track made of a lullaby and white noise was played ambiently. The 3D stimuli were rendered using Blender 2.79b (Blender Foundation), the stimuli were presented using MatLab 2006a (7.2.0.232) and Psychtoolbox 3 3.0.9 (beta).

### EEG recording and analyses

The participants’ electrical brain activity was continuously recorded using a Hydrocel Geodesic Sensor Net (Electrical Geodesic Inc.), consisting of 128 silver-silver chloride electrodes evenly distributed across the scalp (124 electrodes were used). The vertex served as the reference. The electrical potential was amplified with 0.1 to 100 Hz band-pass, digitized at 500 Hz sampling rate and stored for off-line analyses^54^. The raw data were processed offline using NetStation 4.5.1 analysis software (Electrical Geodesic Inc.). Continuous EEG data were high-pass filtered at 0.3 Hz and low-pass filtered at 30 Hz using digital elliptical filtering^54^. They were then segmented in epochs from 300 ms before the tactile stimulus onset until 1300 ms after it and baseline-corrected to the average amplitude of the 100 ms interval preceding the tactile stimulus onset. Epochs containing movement artefacts were visually detected and rejected, as well as epochs with more than 12 bad electrodes^54^. Bad electrodes (if less than 12) were interpolated on a trial-by-trial basis using spherical interpolation of neighbouring channel values.

Video recordings of the experimental session were made and then observationally coded offline to identify any trial in which: i) the participant was not looking at the screen, ii) the participant was looking at the moving stimulus rather than the attention getter, iii) the participant’s hands were not in the correct position. If found, these trials were excluded from further analyses. Artefact free data were rereferenced to the average potential over the scalp, then individual averages were calculated. The average numbers of artefact free trials considered for the analyses for each age group, condition and trial type are summarised in Table 5. The relatively small number of trials available per condition could be due to the high number of different conditions in which the infants participated (N = 4) and is not unusual in infancy research^54^.

**Table 5.** Mean number of artefact-free trials considered for analyses for each age group, condition and trial type (parenthetical values are SDs).

|  | **Approaching**<br>**Touch** | **Approaching**<br>**No-Touch** | **Receding**<br>**Touch** | **Receding No-**<br>**Touch** |
| --- | --- | --- | --- | --- |
| 4-month-olds | 10.05 (3.15) | 10.20 (3.72) | 9.40 (3.12) | 9.75 (3.60) |
| 8-month-olds | 9.85 (2.13) | 9.25 (2.67) | 9.50 (2.65) | 9.90 (2.95) |

We were interested in analysing the SEP responses originating from sites close to somatosensory areas, as our aim was to investigate infants’ brain responses to tactile stimuli preceded by unattended approaching or receding visual stimuli. In order to make sure that the analysed waveforms were not affected by the visual components of the stimulation, we computed and compared the difference waveforms obtained subtracting, for each visual motion condition (Approaching vs Receding), the response recorded in the No-Touch trials from the response recorded in the Touch trials (see Design). The difference waves obtained represented the response to the tactile stimulus itself, given that visual components, common across the two types of trials, were removed by the subtraction.

To identify the clusters of electrodes to be used for the analyses, first of all we averaged the difference waves. Then, we inspected the averaged topographic maps representing the scalp distribution of the electrical activity^47^, which confirmed the presence of hotspots in the regions surrounding CP3 and CP4 in the 10-20 system^55,56^. Next, we visually inspected the averaged recordings from the electrodes within these areas to isolate, for each hemisphere, the cluster of electrodes showing the most pronounced SEP components^51^. The electrodes chosen for the analyses were: for the 4-month-old group, 36, 41, 42 (left hemisphere) and 93, 103, 104 (right hemisphere); for the 8-month-old group, 41, 46 47 (left hemisphere) and 98, 102, 103 (right hemisphere).

### Statistical information

To investigate any differences in infants’ SEPs in response to tactile stimuli preceded by approaching vs receding motion, first of all we ran a sample-point by sample-point analysis, using the Monte Carlo simulation method^46^. This method allowed us to identify the time course of statistically reliable modulations of the SEPs, correcting for the auto-correlation of consecutive sample points (i.e. 2 ms intervals). This analysis simulated 1000 randomly generated datasets having the same level of auto-correlation and the same number or participants and samples points of the observed data, and calculated the shortest length of consecutive significant values reliably significant with 95% probability, i.e., not generated by chance by the statistical dependence of consecutive time points^46^.

Then, we identified by visual inspection any significant components present within the time window deemed as significant by the simulation. The exact time window for each component was determined by averaging the waveforms across participants and conditions, in order to avoid biased measurements^47^. We calculated and compared the mean individual amplitude of the responses within these components using paired-planned two-tailed t-tests. Effect sizes (Cohen’s *d_z_*) were calculated dividing the mean of the differences between the mean individual amplitude values by the standard deviation of the same differences. All reported statistical tests for both age groups were conducted on the full final sample, N = 20.

Then, for exploratory purposes, we ran a number of LMMs. The models were fitted using the R software^57^, specifically the “lme4” and “lmerTest” packages for LMMs^58,59^. They were compared pairwise using Likelihood Ratio Tests (LRTs) and the best fitting mode was fitted by Restricted Maximum Likelihood (REML). The t-tests in the model summary use Satterthwaite’s method.

## Data availability

The datasets generated and/or analysed during the current study are available in the University of Birmingham eData repository^60^ and can be retrieved from https://doi.org/10.25500/edata.bham.00000447. The scripts and datasets used to perform the analyses reported in this manuscript are available online on the Open Science Framework website^61^, and can be retrieved from https://osf.io/jg7xf/.

## Acknowledgements

We are grateful to our participants and their parents, for their invaluable contribution. This research was supported by a British Academy/Leverhulme Trust grant (SG170901) awarded to G.O. and A.J.B. and a Leverhulme Trust Early Career Fellowship (ECF-2019-563) awarded to G.O.

## Author contributions

G.O. conceived the study, designed it, collected the data, undertook data processing and analyses and wrote the report. I.P. collected the data, and undertook some data processing. A.J.B. and J.v.V. contributed to the design of the study and the interpretation of the findings. A.J.B. helped writing the report.

## Additional Information

Competing Interests: The authors declare no competing interests.

